# A novel, ultrasensitive approach for quantitative carbohydrate composition and linkage analysis using LC-ESI ion trap tandem mass spectrometry

**DOI:** 10.1101/853036

**Authors:** Kathirvel Alagesan, Daniel Varon Silva, Peter H Seeberger, Daniel Kolarich

## Abstract

Glycan identification and characterisation is essential to correlate glycoconjugate structure to biological function. The structural assignment of carbohydrates is often based on MS composition analyses and knowledge on well-studied glycosylation pathways. Nevertheless, many monosaccharide building blocks are indistinguishable by mass alone and detailed linkage information is also not easily obtained by MS/MS analyses, in particular when organisms are studied where the glycosylation pathways are less well defined. Here, we present a novel, simple and sensitive method using Reversed Phase (RP) – Liquid Chromatography Electrospray ionisation tandem mass spectrometry (LC–ESI-MS/MS) for unambiguous identification and linkage determination of monosaccharides including N-acetylneuraminic acids. Sequential permethylation and reductive amination steps are employed prior and after acid hydrolysis to enable separation and differentiation of the various monosaccharides and their respective linkage positions. The well-established, monosaccharide specific methylation patterns allowed for the identification of the various derivatised monosaccharide alditols based on their retention time and tandem mass spectrometry fingerprint. Absolute quantitation can also be accomplished by including a set of internal standards, thus simultaneously providing qualitative and quantitative information on the monosaccharide residues present.

## INTRODUCTION

Carbohydrates and their diverse conjugates are crucial for life as we know it, both as an energy source and as key structural and functional components of cell walls and membranes. In-depth knowledge of the primary sequence of oligo- and polysaccharides is vital to understand the biology of glycoconjugates (1). The *de novo* comprehensive structural analyses of complex carbohydrates are labour-intensive and time-consuming, requiring orthogonal methods to identify (i) the constituent monosaccharides, ii) their sequence including branching points, iii) glycosidic linkage configuration, iv) position of glycosidic linkage and v) elucidation of anomeric configurations (2, 3). Next to interaction (lectins/antibodies) and NMR based approaches, various LC and LC-MS based methods are available for glycan profiling that provide different aspects of this information either directly on intact glycan structures or after hydrolysis of these into their monosaccharide building blocks (4). Glycomics analyses have significantly advanced in the past decades and are now commonly applied in a rapid-throughput manner (5, 6).

LC, MS or LC-MS based glycan composition and structure assignment inherently relies upon pre-existing knowledge on glycan biosynthetic pathways and identities of monosaccharide residues. Hence, the lack of this information severely limits composition/structure assignment when studying glycoconjugates in less characterized species. Therefore reliable, sensitive, and selective methods enabling the determination of monosaccharide identities and their linkages are highly relevant. While MS has developed to be one of the most powerful technologies for structure analyses, it struggles to differentiate isobaric monosaccharide residues and is not always sufficient to inform on the linkage between monosaccharide building blocks.

Among the traditional methods for monosaccharide composition and linkage analysis, gas chromatography coupled to mass spectrometry (GC-MS) has been the method of choice for decades to unravel monosaccharide identity and linkage information within a single analytical experiment (7). This method is based on the conversion of monosaccharides into partially methylated alditol acetals (PMAAs) that are the products of a series of derivatization steps: permethylation, acid hydrolysis, reduction, and following acetylation of the partially methylated sugar alditols. Different monosaccharide residues and their linkages can be easily identified based upon their GC retention time and characteristic fragmentation patterns (8-10) This method requires a dedicated GC-MS instrument, which is nowadays hardly available for routine analyses in glycomics and glycoproteomics laboratories.

Alternative methods such as high-pH anion-exchange chromatography with pulsed amperometric detection (HPAEC-PAD), capillary electrophoresis (CE), and high-performance liquid chromatography (HPLC) separation coupled with UV/fluorescence detection are also commonly used for monosaccharide analyses (7). The use of specific exoglycosidases in combination with these techniques allows elucidation of oligosaccharide sequence, monosaccharide identity, anomeric configuration, and linkage due to the well defined, high substrate specificity of many of these enzymes (11, 12). This method, however, requires a vast array of enzymes and sufficient amounts of material also a considerable amount of time and appropriate controls to ensure proper enzyme activity to achieve complete and reliable sequencing. Also, this approach fails when suitable exoglycosidases are unavailable.

With the exception of HPAEC-PAD (13) derivatization of monosaccharides with a UV-absorbing or fluorescent tag is necessary to improve monosaccharide HPLC separation and facilitate their detection (14). Several alternative methods have been reported using LC-MS for the analysis of monosaccharides. There, a post-column addition of Na^+^, Cs^+^ and NH_4_^+^ or I^−^ and Cl^−^ ions, respectively, have been used to generate positively (or negatively) charged molecules (15-20). A few LC-MS based approaches have been reported for underivatized monosaccharides (21-23), but many of these approaches lack sensitivity or are unable to provide valuable linkage information. Not until very recently an LC-MS based method was reported for the rapid and simultaneous relative quantitation of glycosidic linkages for oligosaccharide and polysaccharide characterization, however, lacked baseline separation of 6-galactose and 6-glucose; 6-mannose and 4-galactose (24). Here, a novel, highly sensitive nanoLC-ESI ion trap tandem mass spectrometry-based method for monosaccharide composition and linkage analysis is presented that also allows differentiation of isobaric monosaccharides. The method enables the simultaneous labelling of both, neutral and acidic monosaccharides using aniline, the relative quantitation of the different monosaccharides within a given sample and also informs on the branching points.

## MATERIALS AND METHODS

### Reagents

- L(−)Fucose (Sigma-Aldrich, cat. no. F-2252)
- D(+)Xylose (Sigma-Aldrich, cat. no.X1500)
- D(+)Mannose (Sigma-Aldrich, cat. no. M-4625)
- D(+)Galactose (Sigma-Aldrich, cat. no. G-0750)
- D(+)Glucose (Sigma-Aldrich, cat. no. G-8270)
- N-Acetyl-D-glucosamine (Sigma-Aldrich, cat. no. A8625)
- N-Acetyl-D-galactosamine (Sigma-Aldrich, cat. no. A2795)
- N-Acetyl-D-mannosamine (Sigma-Aldrich, cat. no. A8176)
- N-Acetylneuraminic acid (ACROS Organics cat. no. 227040250)
- DMSO (ACROS Organics, cat. no. 127790010)
- CH_3_I (Sigma-Aldrich, cat. no. 67692)
- CD_3_I (Sigma-Aldrich, cat. no. 176036)
- NaOH (Sigma-Aldrich, cat. no. S5881)
- Glacial acetic acid (Merck, cat. no. 100066)
- TFA (Merk, 1081780050)
- Sodium cyanoborohydride (Sigma-Aldrich, cat. no. 156159)
- DCM (Merck, cat. no. 6048)
- Chloroform (Merck, cat. no. 102445)
- *N*-glycan standards (Dextra Reading, UK)
- ZipTips (Millipore)

Water was used after purification with a Milli Q-8 direct system (Merck 119 KGaA, Darmstadt, Germany). Stock solutions with a concentration of 1 mM of each monosaccharide (Glc, Gal, Man, GlcNAc, ManNAc, GalNAc, Xyl, Fuc, Neu5Ac) were prepared using MilliQ-8 water.

### General procedures for carbohydrate derivatisation

Standard monosaccharides were labelled by reductive amination using 2-aminobenzamide (2-AB) as described earlier (25). Briefly, labelling reagent was prepared by dissolving 0.35 M of 2-AB and 1 M sodium cyanoborohydride in 30% acetic acid in DMSO. About 5 mg of monosaccharides were dissolved in 100 μL of the labelling reagent, and the reaction mixture was incubated at 65°C for 120 min. Excess reagent was removed as described by Pabst *et al.* previously (26).

Monosaccharide permethylation was performed according to the procedure described previously by Ciucanu & Kerek (27) with minor modifications as described by Ciucanu I & Costello CE (28). Briefly, the glycan sample (approx. 5 mg of standard monosaccharides) was dissolved in 1000 μL of DMSO by gentle vortexing and 50 mg of finely powdered NaOH was added. The mixture was sonicated for 10 min at room temperature and incubated for 30 min with occasional shaking. Subsequently, methyl iodide (100 μL) was added and the sample was sonicated for another 10 min. The reaction was terminated by the addition of 1000 μL water. Permethylated monosaccharides were extracted using 1000 μL of chloroform. The chloroform layer was washed with equal volumes of water for at least 3 times to remove any residual salts. The organic phase containing the permethylated monosaccharides was dried *in vacuo* and reconstituted in 20% (v/v) aqueous acetonitrile.

A detailed step-by-step procedure for monosaccharide linkage and compositional analysis is provided in supplementary section.

### ESI-MS/MS and LC-ESI-MS/MS analysis of the derivatised monosaccharides

All mass spectrometric analyses were performed on an amaZon ETD Speed ion trap (Bruker, Bremen, Germany) coupled to an Ultimate 3000 UHPLC system (Dionex, Part of Thermo Fisher Scientific, Germany).

Differently derivatized monosaccharide standards were dissolved in 30% acetonitrile containing 0.1% formic acid (FA) at the calculated concentration of 50 pmol/μL. All the samples were directly infused into the ESI-source for MS analysis using a syringe pump at a flow rate of 1 μL/min. The instrument was set up to perform CID fragmentation in positive mode on the selected precursors. Precursor ions were selected manually with the isolation width of ±1 Da, and the fragmentation energy was increased manually to determine the optimal fragmentation conditions. The data was recorded in the instrument’s “enhanced mode resolution”. Specific instrumental operational parameters used in the present investigation are listed in the Supplementary Table S1.

On the nanoLC system the samples were injected (analyte concentration: 50 pmol) onto the precolumn in 100% solvent A (0.1% formic acid) at the flow rate of 6 µL/min. Unbound components were washed off from the column for 5 min with buffer A. The analytical column was equilibrated in 2% solvent B (acetonitrile with 0.1% formic acid) and a linear gradient was applied using an increasing solvent B concentration at the flow rate of 300 nL/min as follows: steep increase of buffer B from 1 to 13% (from 5 min to 8 min), followed by a slow increase of buffer B from 13% to 30% (8 to 70 min) and then a steep increase to 90% (70 to 75 min). The column was held at 90% B for 5 min before re-equilibrating the analytical column in 1% solvent B. Meanwhile the precolumn was re-equilibrated in 100% solvent A prior injection of the next sample.

## RESULTS AND DISCUSSION

### Rationale and Method Development

Monosaccharide analysis is critical to identify the building blocks of complex carbohydrates. In-depth structural analysis of carbohydrates, in particular when derived from less well studied organisms, remains a daunting task due to the lack of sensitive LC-MS based analytical methods that allow differentiation of monosaccharide isomers and their linkage(s). We aimed to overcome these limitations by developing an easily adaptable and sensitive nanoLC-ESI MS/MS-based workflow for this purpose. The method employs sequential derivatization steps and strong acid hydrolysis at elevated temperatures, followed by the analysis of the labelled monosaccharide by a standard proteomics setup using RP-nanoLC-ESI-MS/MS.

To achieve the highest possible sensitivity, various derivatization and labelling steps were first optimized using standard monosaccharides (***Supplementary Figure 1***). Next, the fragmentation behaviour and chromatographic properties of these derivatized monosaccharides (*Table 1*) were evaluated using various stationary phases (reverse phase C18, porous graphitized carbon, PGC) and the chromatographic conditions were optimized to separate the various monosaccharide stereoisomers.

**Table 1:** Protonated [M+H]^+^ mass values of the differently derivatized monosaccharides analysed in the course of this work

| Compound | Mono-saccharide | Monoisotopic mass | 2-AB | 2-AB Me | Aniline | Aniline-Me |
| --- | --- | --- | --- | --- | --- | --- |
| Hexose | Glc, Gal, Man | 181.06 | 301.13 | 413.26 | 258.12 | 342.22 |
| HexNAc | GlcNAc, GalNAc, ManNAc | 222.08 | 342.16 | 454.29 | 299.14 | 383.32 |
| Deoxyhexose | Fuc | 165.06 | 285.14 | 383.25 | 242.12 | 312.12 |
| Pentose | Xyl | 151.13 | 271.12 | 369.23 | 228.11 | 298.12 |
| N-Acetyl neuraminic acid | NeuAc | 310.10 | - | - | 387.16 | 498.13 |
| 2-AB – 2 aminobenzamide labelled monosaccharide<br>2-AB-Me – 2-aminobenzamide labelled, permethylated monosaccharide<br>Aniline-Me – Aniline labelled, permethylated monosaccharide |  |  |  |  |  |  |

### Stereochemistry defines the fragmentation of 2-AB labeled, permethylated monosaccharides

Offline MS analysis of the 2-AB labelled, permethylated (2-AB-Me) Hex and HexNAc molecules resulted in the preferential formation of protonated [M+H]^+^ ions, whereas Fuc and Xyl were detected as both, protonated and sodiated ion species. For these initial MS/MS analyses the [M+H]^+^ ions were selected, and the fragmentation energies manually varied to determine the optimal fragmentation conditions that provided maximum information. The MS/MS spectra of all the three investigated Hexoses produced comparable fragmentation patterns, but showed distinct fragmentation signatures in the fragment ions intensities that were specific for each compound analysed (*Figure 1*). A successive loss of CH_3_OH from the *m/z* 413.3 precursor ion produced the observed *m/z* 381.2; 349.2; 317.2; 285.2 product ions. The respective monosaccharides’ stereochemistry attributed to this phenomenon by dictating the fragmentation route.

**Figure 1:**
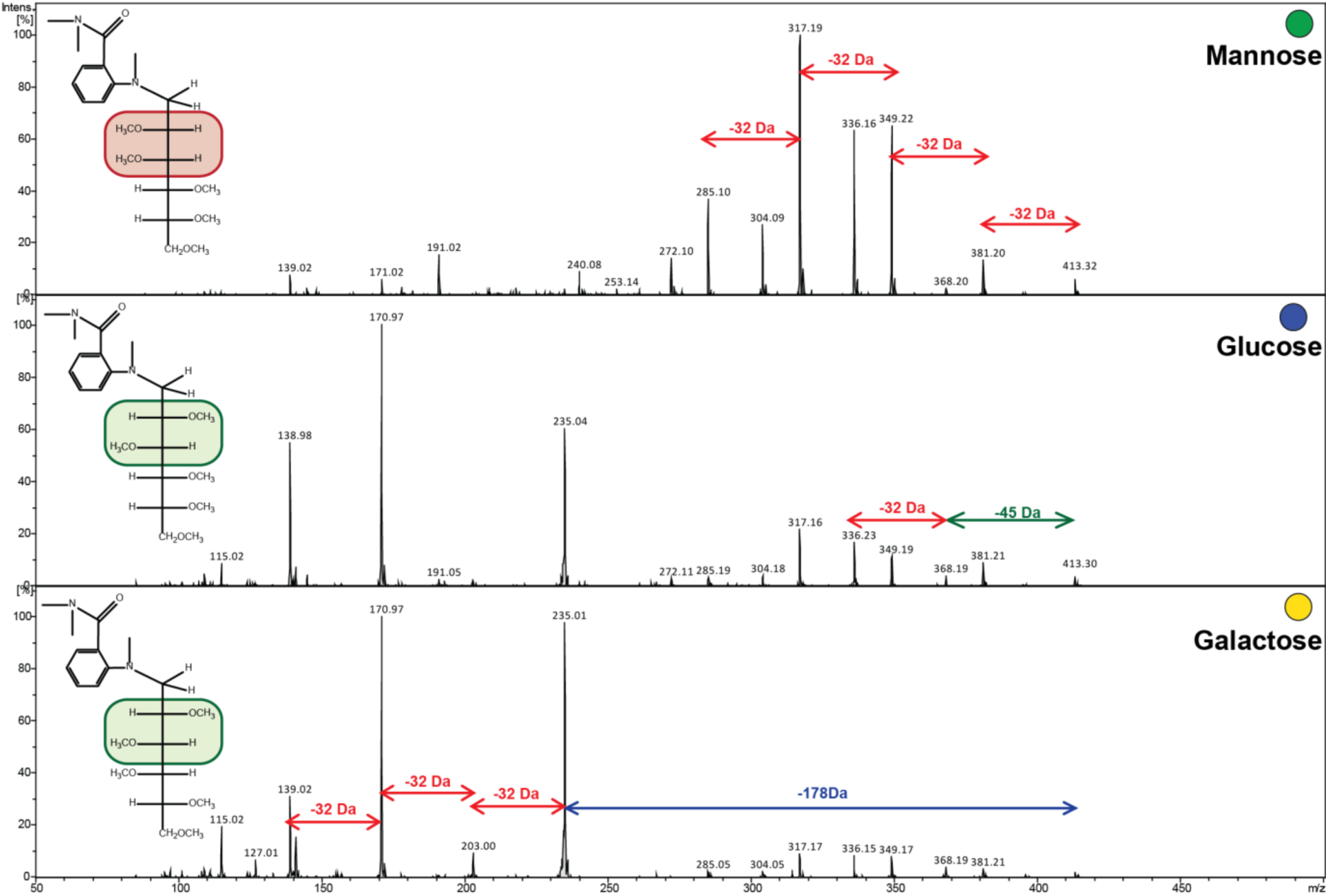
Fragmentation pattern of 2-AB-Me hexoses of the protonated precursor ion *m/z* 413.3. Product ions observed at *m/z* 381.2; 349.2; 317.2; 285.2 are the result of a successive loss of CH_3_OH from *m/z* 413.3. The cleavage of the C–N bond between the sugar and fluorescent tag is defined by the sugar stereochemistry.

In comparison to the hexoses the MS/MS spectra of 2-AB-Me GlcNAc were rather simple. The product ion at *m/z* 276.2 deriving from the *m/z* 454.2 precursor followed a similar fragmentation as shown in ***Supplementary Figure 2***. In contrast to the 2-AB-Me hexoses, the MS/MS spectra of all three analysed 2-AB-Me HexNAc’s were fairly similar and less informative on the respective stereochemistry. The observed fragmentation spectra, however, still provided a sufficient number of fragment ions to differentiate these compounds from other low-molecular-weight compounds, and their different retention time allowed an easy identification (Figure 3).

The MS/MS spectra obtained for 2-AB-Me fucose and xylose exhibited fragmentation signatures similar to the ones of hexoses with the successive loss of 32 Da from the protonated precursor ion (***Supplementary Figure 3***). The loss of 178.1 Da from the protonated precursor ion was consistent for all investigated 2-AB-Me derivatised monosaccharides (*Figure 2*). The sodiated molecular ions followed the same routes albeit larger fragmentation energies were required to achieve sufficient fragmentation.

**Figure 2:**
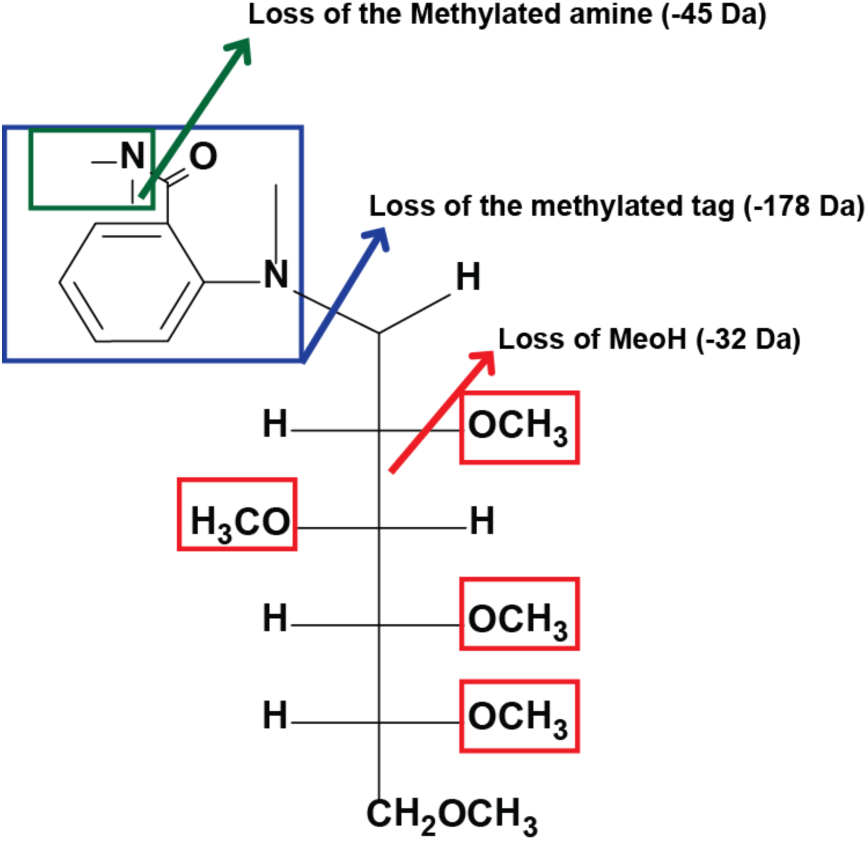
Fragmentation of 2-AB Me derivatised monosaccharides during electrospray ionisation. Primary fragments occur between the C1 atom and the Nitrogen atom of the fluorescent tag, which results in the neutral loss of 178 Da from the precursor. Secondary fragmentation results in the loss of methanol (32 Da) and methylated amine (45 Da) from the precursor mass.

### Chromatographic separation of 2-AB-ME labeled monosaccharides

Most monosaccharides occurring in mammalian glycoconjugates are isobaric stereoisomers (Table 1) that cannot be differentiated by a simple MS analysis unless these isomers are separated by orthogonal means such as liquid chromatography. Different stationary phases were tested for their ability to achieve separation of the different isobaric, 2-AB-Me labelled monosaccharides. PGC is well known for its particular ability to separate native oligosaccharides (29). However, it was unable to provide the baseline separation necessary to deliver single peaks for the various 2-AB-Me hexoses (data not shown). The intended separation was achieved using C18 reversed-phase stationary phases after optimizing gradient and solvent conditions (Figure 3).

**Figure 3:**
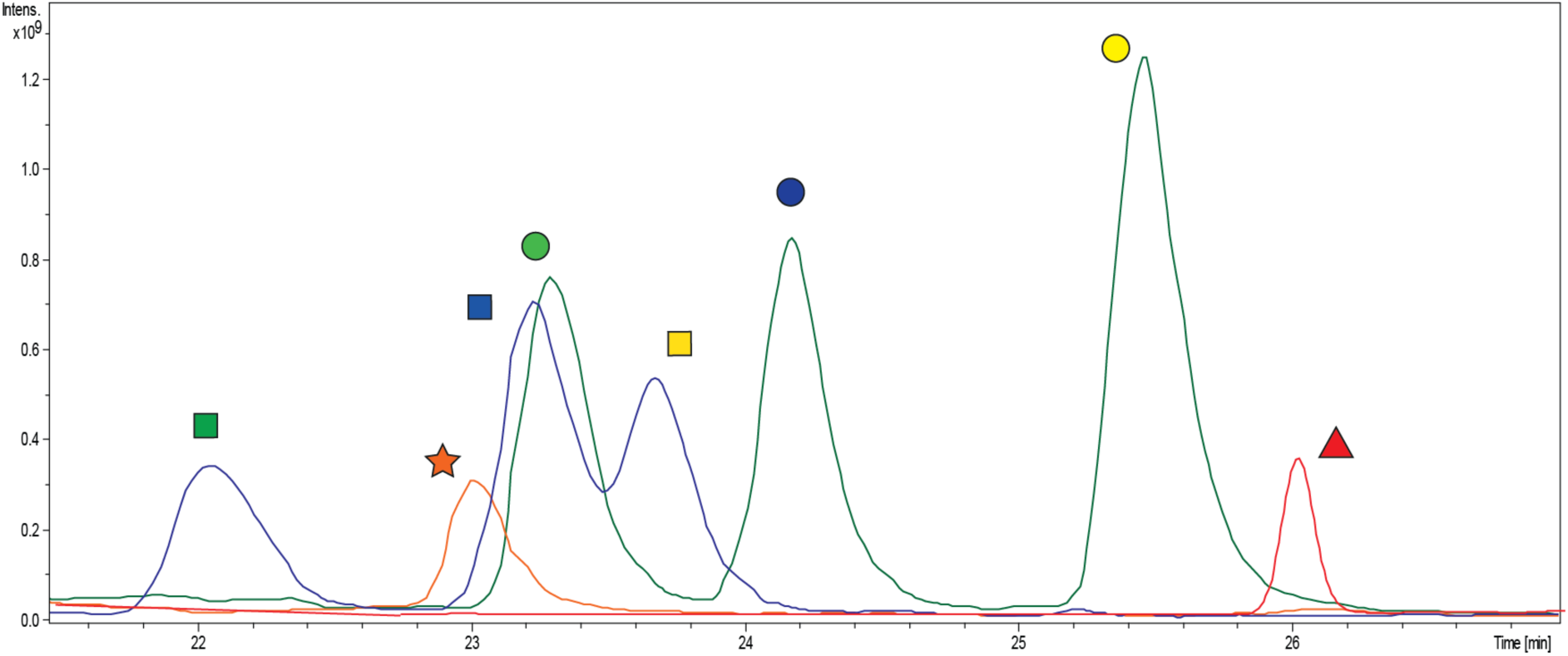
LC-ESI-MS/MS analysis of various 2-AB-Me derivatised monosaccharides separated using RP-C18 chromatography. Calculated 50 pmol of the respective compounds were injected onto the column. Extracted Ion Chromatogram (EIC) traces are shown of various monosaccharides separated under conditions described in material and methods section.

### Evaluation of the linkage analysis strategy using reference oligosaccharides

The successful establishment of an analysis workflow suitable for 2-AB-Me labelled monosaccharides was the basis to further develop the linkage analysis approach (Figure 4). This strategy was based on the principle that first, the reducing ends of the target oligosaccharides were derivatized by reductive amination with 2-AB. In the next step, the free hydroxyl groups were permethylated by CH_3_I, which also resulted in the methylation of all amino groups. These 2-AB-Me modified oligosaccharides were then subjected to acid hydrolysis, followed by a second reductive amination with 2-AB of the newly generated, free reducing ends of the partially methylated monosaccharides. In a second permethylation step, the use of CD_3_I allowed the introduction of a CD_3_ label on all C atoms that were not permethylated during the first permethylation reaction as they were involved in glycosidic linkages. Permethylation using CD_3_ resulted in an additional mass increase of +3 Da at each position previously involved in a glycosidic linkage.

**Figure 4:**
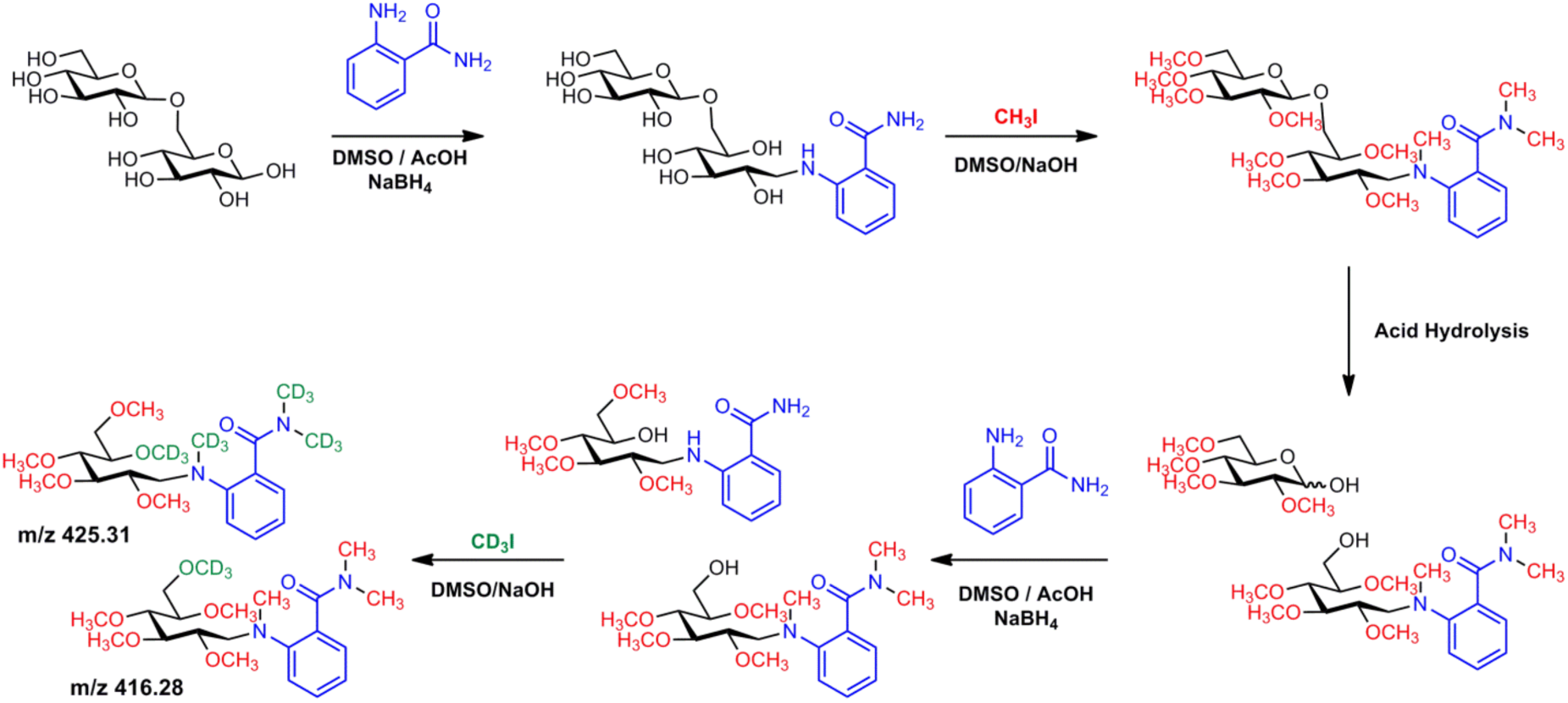
Linkage analysis workflow principle explained using gentiobiose, a *α*1-6 linked glucose disaccharide. Sequential permethylation and reductive amination steps prior and after acid hydrolysis enable monosaccharide identification and linkage characterization based on specific signature MS/MS spectra.

A proof of principle evaluation of this strategy was performed on various reference oligosaccharide standards. Gentiobiose, lactose, trehalose, and maltoheaptose were first tested before the approach was also applied to complex *N*-glycan standards. As demonstrated on the gentiobiose example the reducing end and non-reducing end glucose could be differentiated just by their molecular masses, as these differed as a consequence of this specific multi-step labelling/permethylation strategy (Figures 4 and 5-A). Also, all linkage information could be deduced from the individual fragmentation patterns (Figure 5-B).

2-AB-Me monosaccharides exhibited successive 32 Da losses from the precursor mass (**Fragmentation route 1**, Figure 2). In the presence of CD_3_, however, 35 Da losses were detected at the respective linkage position, providing sufficient information to assign the C-atom involved in oligosaccharide formation. Signals derived from the proposed fragmentation route 1 provided information on the reducing and non-reducing end monosaccharides, respectively (Figure 5-B). In the presence of CD_3_ a slight shift towards earlier retention times was observed when compared to the respective CH_3_ methylated monosaccharides (Figure 5-A). Such retention time shifts have also been described earlier for deuterated peptides and were associated with differences in the size of hydrogen and deuterium atoms and their binding interaction energies to the stationary phase (30). This phenomenon, however, did not alter the base line separation of the various monosaccharide stereoisomers and thus did not affect the here developed linkage and monosaccharide analysis approach. Linkage analysis results obtained from the standard disaccharides trehalose, lactose, and gentiobiose are shown in ***Supplementary Figures 4+5.***

**Figure 5:**
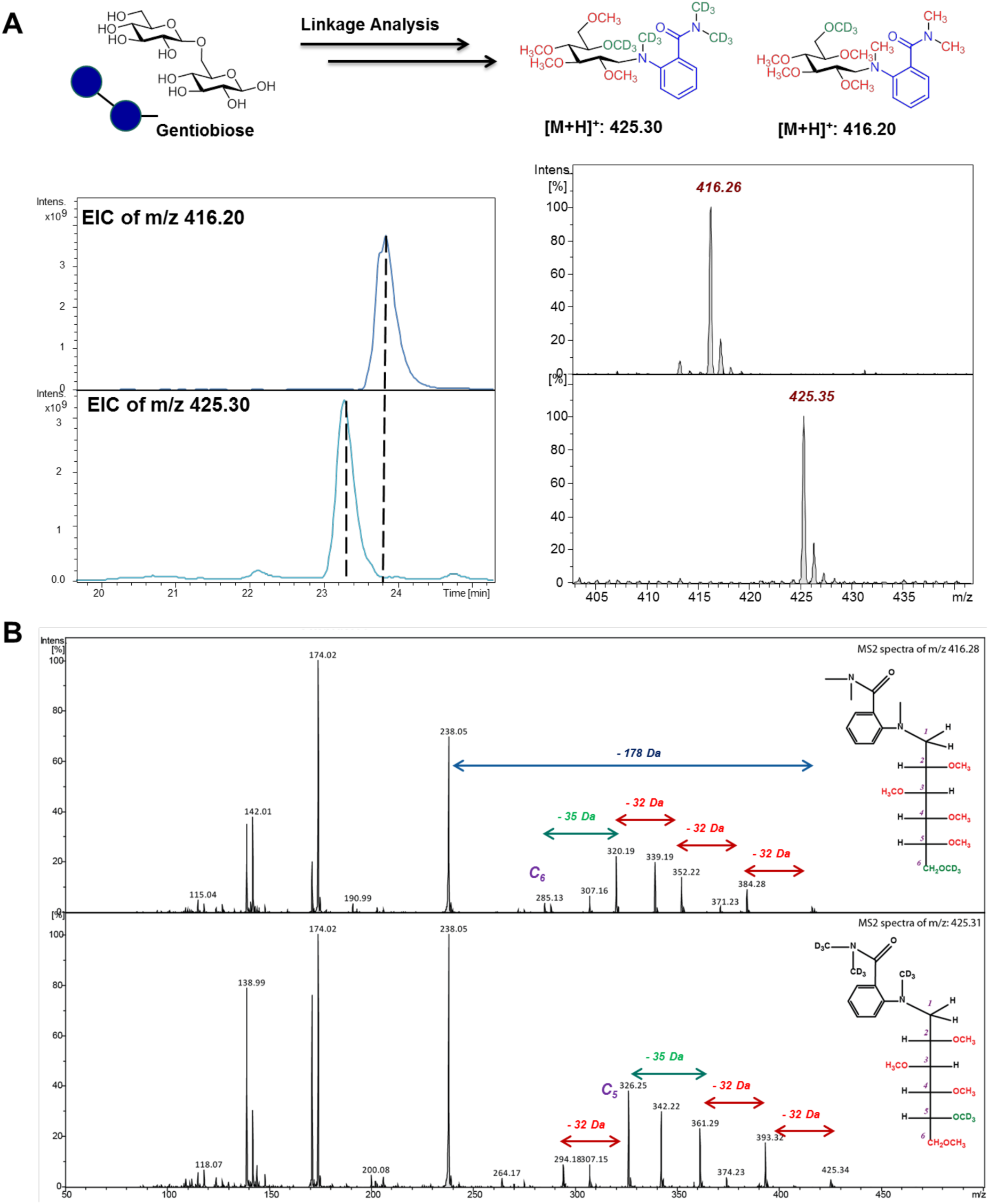
LC-ESI-MS/MS Analysis of 2-AB methylated monosaccharides derived from the standard disaccharide gentiobiose after linkage analysis. (A) simplified scheme of linkage analysis and EIC of the 2-AB-Me derivatized monosaccharides obtained after linkage analysis.(B) MS/MS spectra of the derived 2-AB-Me monosaccharides.

### Method Application to *N*-Glycans

The applicability of this approach to allow monosaccharide analyses from protein released glycans was also tested using a panel of commercially available synthetic *N*-glycan standards (***Supplementary Table 2***). Approximately 200 fmol of the reduced *N*-glycan alditols were subjected to linkage analysis. In the case of the oligomannose type *N*-glycan carrying an additional glucose residue on the 3-arm two different non-reducing end hexoses are obtained, one glucose and two mannose residues per molecule (*m/z* 425.35). Also, five mannose residues that are extended on the C2 position (*m/z* 428.34) and two mannose residues that are extended on the C3 and C6 position (*m/z* 431.34) will derive from this analysis per *N*-glycan molecule (Figure 6). The linkage analysis results for the other standard *N*-glycan is shown in Supplementary Figure 6. This exemplifies that the here developed technique can be easily applied to obtain basic linkage and monosaccharide identity information even from sub-picomol amounts of starting material.

**Figure 6:**
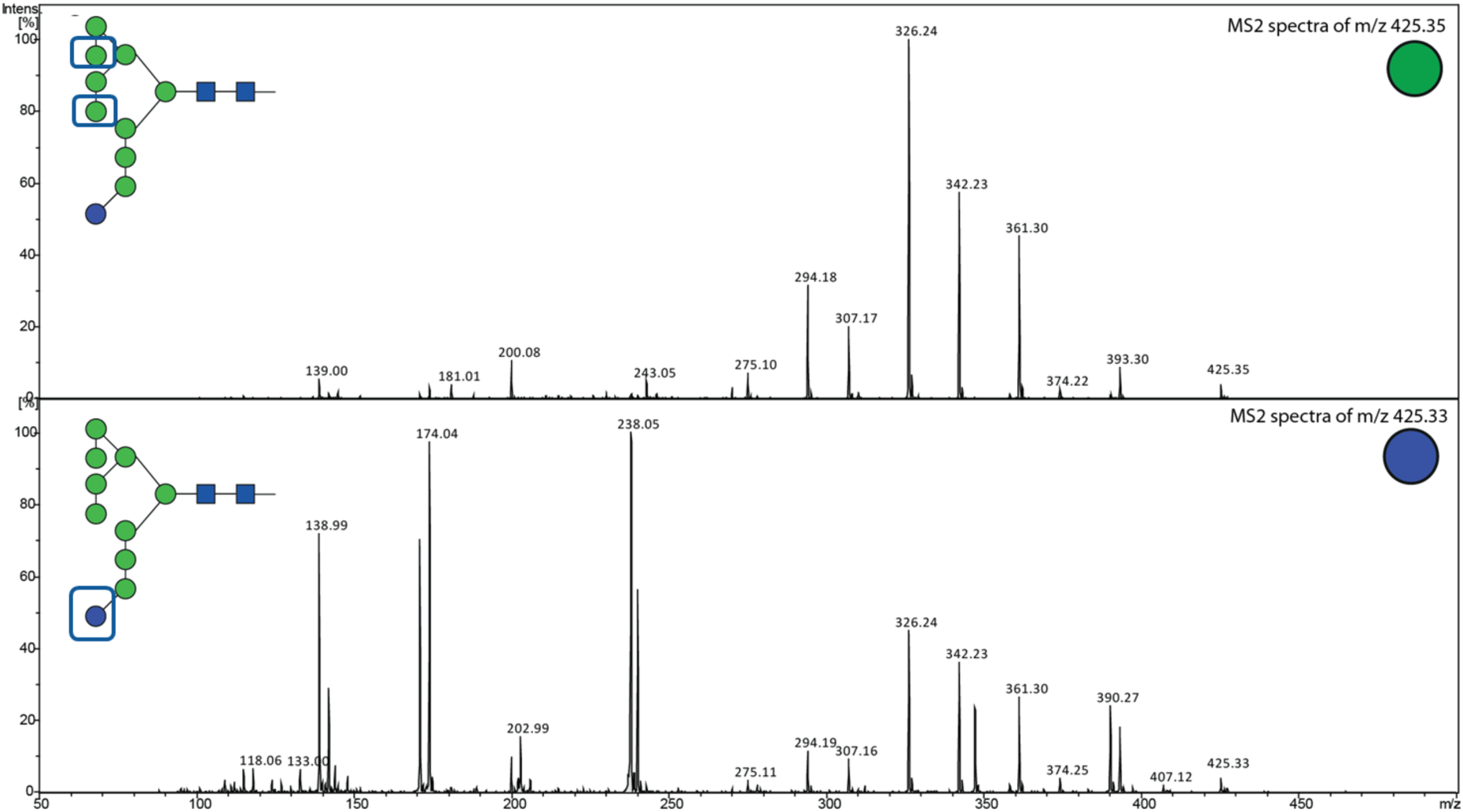
Fragment spectra of terminal mannose and glucose monosaccharides present in the Man9-Glc1 *N*-glycan. Linkage positions of terminal monosaccharides were identified based on the molecular mass, order of elution and the signature fragmentation pattern of 2-AB-Me monosaccharides.

### Quantitation

Any quantitative information on the monosaccharides present in a sample could be determined by spiking in known quantities of pure monosaccharides as an internal reference compounds. The principal feasibility of this approach was evaluated using a maltoheptaose standard that was spiked with a known amount of a monosaccharide mix and subjected to linkage analysis (Figure 7).

**Figure 7:**
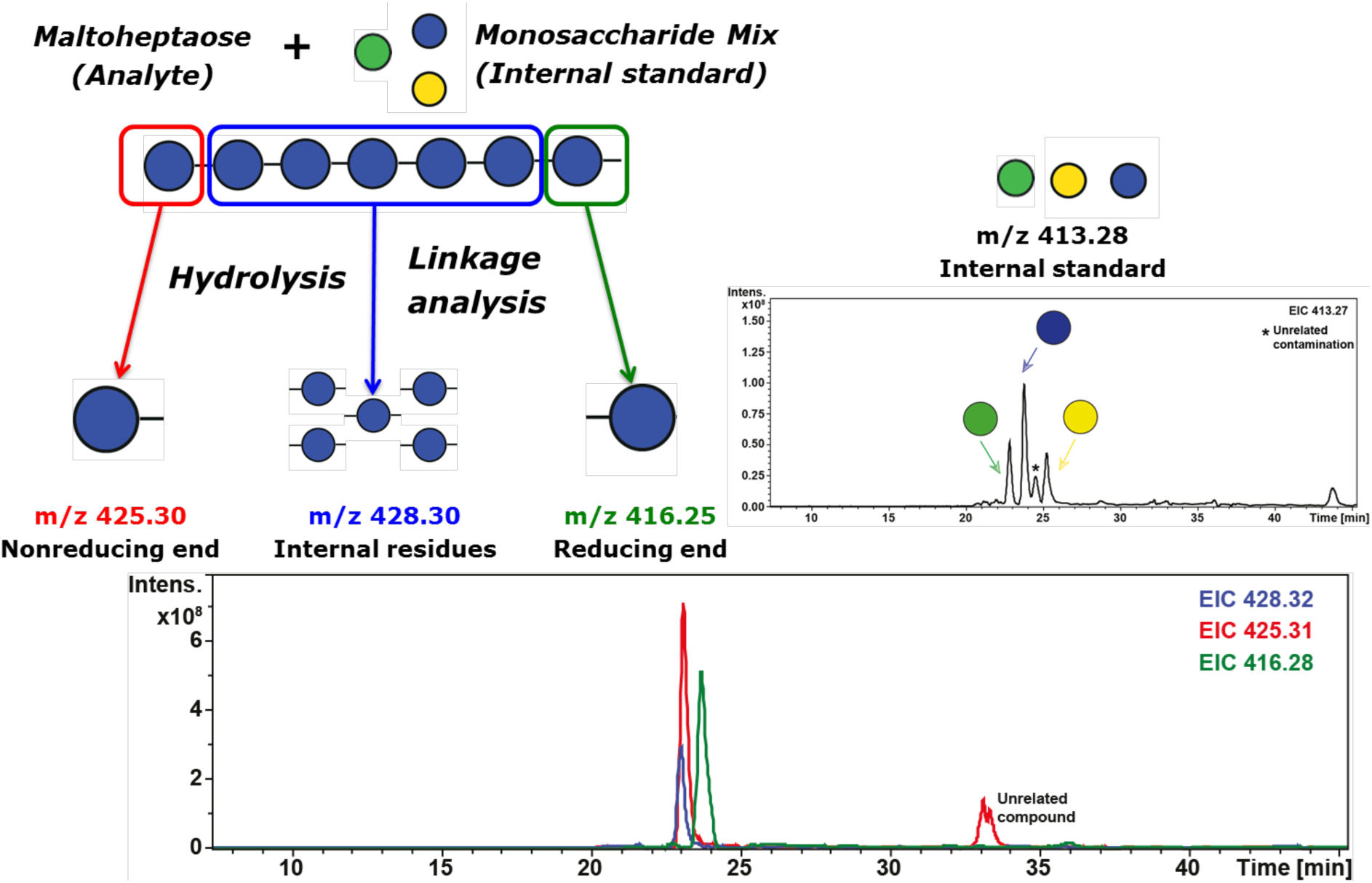
Quantification using internal standards. Known quantities of the standard monosaccharide were mixed with the analyte and subjected to monosaccharide linkage analysis. The presence of CD_3_ methylation present in the analyte derived monosaccharides could be used to discriminate analyte from standards monosaccharides, as the latter only contained CH_3_ methylation. The added monosaccharide served both as internal quantitation standard and for compound identification during the analysis. MS/MS spectra of the 2-AB-Me labelled monosaccharide derived from the analyte maltoheptaose are shown in Supplementary Figure 7.

### Including sialic acids in the monosaccharide analysis equation

The developed approach provided the ability to identify the different monosaccharides and their linkage(s), except for sialic acids as these cannot be labelled by standard reductive amination using 2-AB. DMB (1,2-diamino-4,5 methylenedioxybenzene.2HCl) is a common labelling reagent for sialic acids, which forms a covalent bond via amination cyclisation reaction (31), but this reagent cannot be used to label other aldo-monosaccharides. Searching for a more universally applicable labelling reagent that is suited for both, keto and aldo-monosaccharides, aniline was evaluated due to its high nucleophilic properties in comparison to 2-AB. With aniline aldo-and keto monosaccharides could successfully be labelled on the reducing end. The fragmentation behaviour of all analysed aniline-Me labelled monosaccharides was also similar to that of the 2-AB-Me ones (Supplementary Figure 8). The consolidated fragmentation pattern of aniline-Me monosaccharide is shown in Supplementary Figure 9.

The separation properties of the aniline-Me derivatised monosaccharides on RP-C18 chromatography were also comparable and baseline separation could easily be achieved for isobaric monosaccharides (Figure 8). Interestingly, we observed that the aniline-Me derivatised *N-*Acetylneuraminic acid eluted in two peaks. Depending on the protonation during the reductive amination, aniline reacts with the C2 atom of the *N-* Acetylneuraminic acid to form both the R and S isomers. These isomers could be separated by this approach (Figure 9).

**Figure 8:**
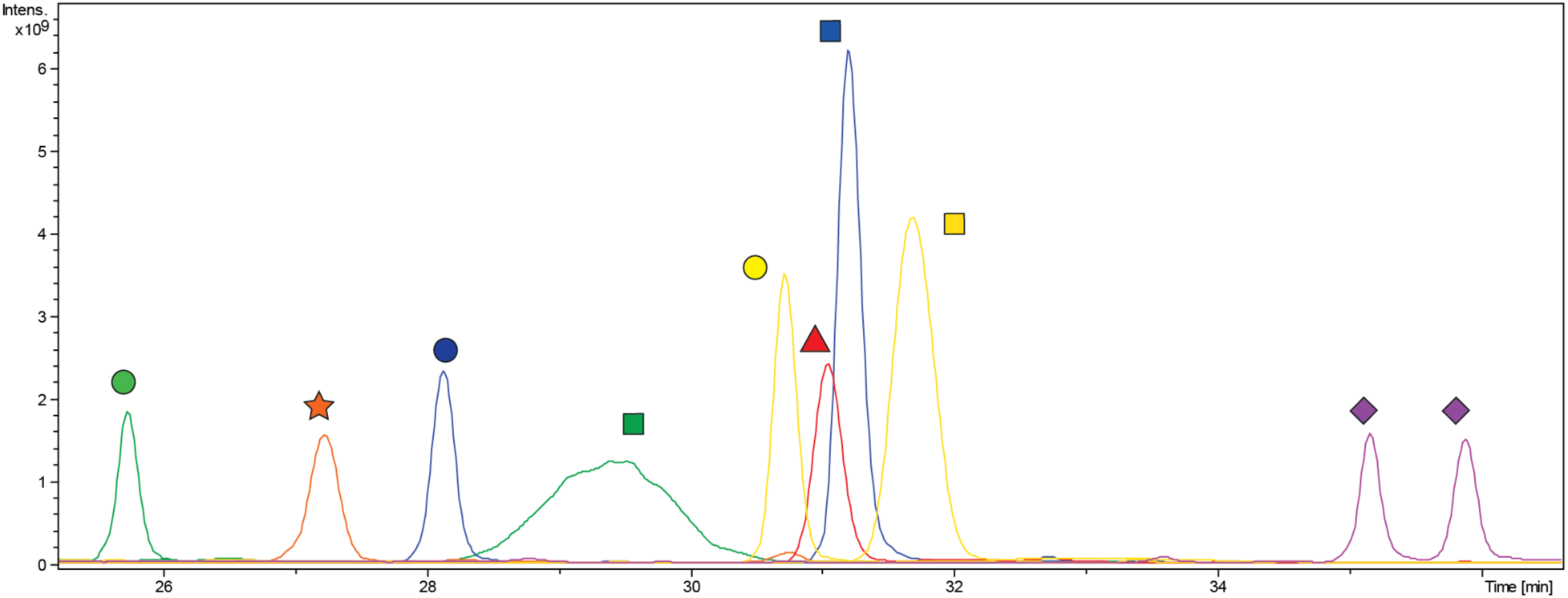
LC-ESI-MS/MS analysis of various Aniline-Me derivatization monosaccharides separated using RP-C18 chromatography. Calculated 50 pmol of the respective compounds were injected onto the column. Extracted Ion Chromatogram (EIC) traces of various monosaccharides separated under optimised conditions are overlaid.

**Figure 9:**
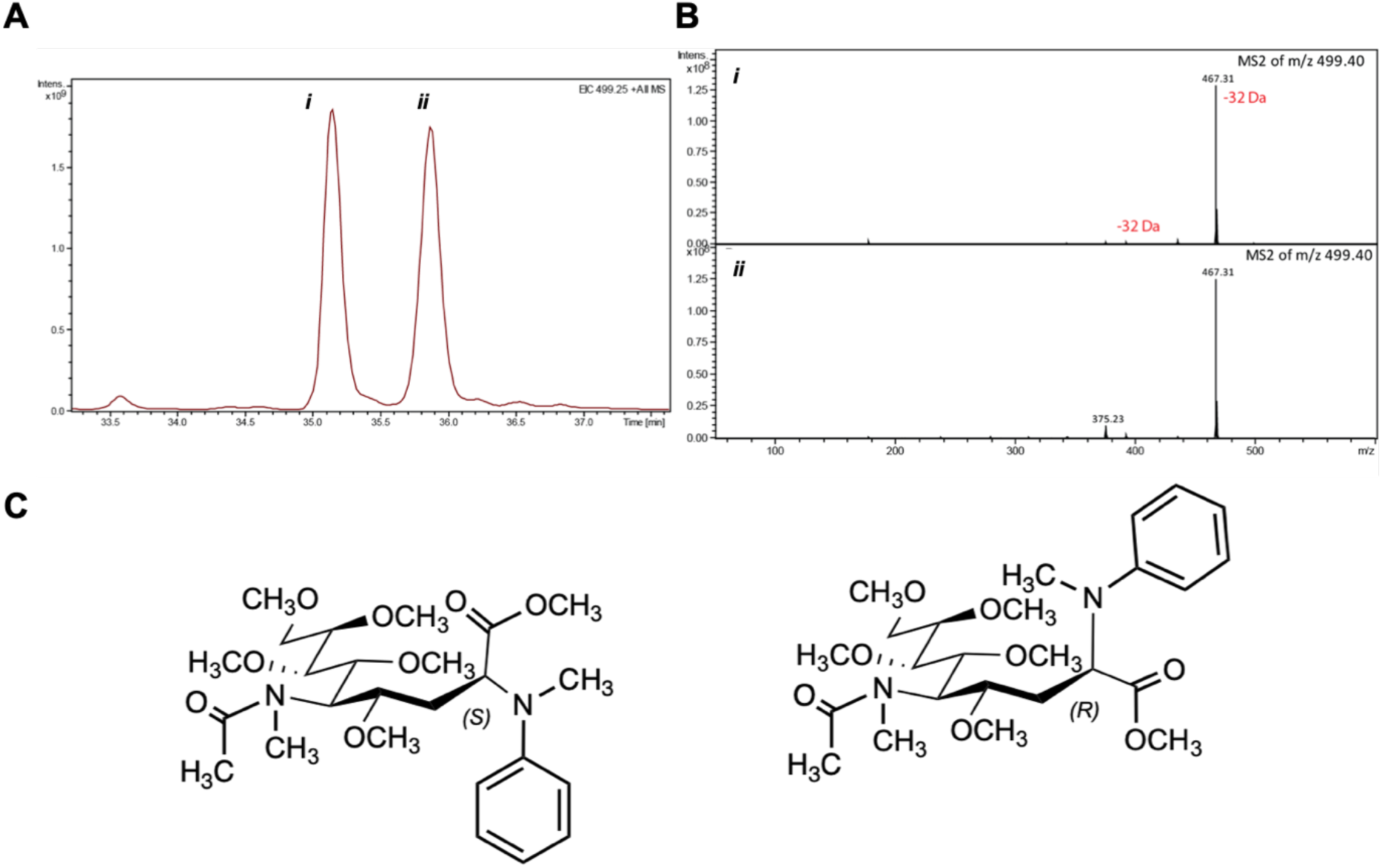
*N*-Acetylneuraminic acid labelling with aniline. (A) EIC-trace of Aniline-Me derivatised *N*-acetylneuraminic acid shows elution of two peaks at different time points. (B) MS/MS spectra of *m/z* 499.40 eluting at the two positions having identical fragmentation spectra. Here we observed the characteristic 32 Da loss from the precursor mass for the aniline-Me derivatised monosaccharides. (C) Chemical structure of R and S isomers of aniline-Me *N*-acetylneuraminic acid resolved by RP chromatography.

In addition, the suitability of aniline as derivatization agent for the linkage analysis was tested using the standard disaccharides gentiobiose and trehalose (Supplementary Figure 10). These results indicated that both aniline, as well as 2-AB, can be employed for linkage analysis providing base line separation of various stereoisomers. Nevertheless, aniline provides an additional advantage of being a universal derivatization agent for both, aldo and keto-sugars.

One point of concern in the current workflow for linkage analysis, however, is that we were unable to detect any *N-*acetylhexosamines (or their respective hexosamine counterparts) in the acid hydrolysis samples due to reasons beyond our explanation. However, the developed approach is still a valuable tool in providing glycan branching point information for each arm from a minimal sample amount.

## CONCLUSION

The work presented here provides a novel, sensitive LC-ESI-MS/MS-based workflow for monosaccharide composition and linkage determination. While nowadays monosaccharide composition and linkage analyses are hardly performed when mammalian glycoconjugates are being characterised, the identification of the building blocks forming glycoconjugates remains vital when studying glycoconjugates from organisms with unknown glycosylation pathways, such as most prokaryotes. While traditional methods used for compositional and/or linkage monosaccharide analysis such as HPAEC-PAD or GC-MS require dedicated instrumentation, we show that similar outcomes can be obtained using a standard nanoLC-ESI MS/MS setup that can be adapted to identify the most commonly occurring monosaccharides including *N*-acetylneuraminic acids. Monosaccharides could be identified after sequential derivatization using the power of C18-RP chromatographic separation combined with mass spectrometric detection. Simultanously, also valuable information on the respective linkages between the monosaccharides was obtained. The reliable limit of detection was determined to be <500 fmol for all analysed monosaccharides, thus providing sufficient sensitivity to successfully analyse as low as 10-20 pmol of initial analyte for a simple compositional monosaccharide profiling. A comprehensive linkage analysis required ≥200 pmol initial analyte, as it requires multiple derivatisation steps. This approach provides one important step forward towards a sensitive, nanoLC ESI MS/MS based approach for the unambiguous assignment of monosaccharide identity and linkage.

## ACKNOWLEDGEMENTS

We thank the *Beilstein-Institut* for supporting KA with a PhD scholarship and the Max Planck Society for the generous financial support. DVS thank the RIKEN-Max Planck Joint Center for Systems Chemical Biology fo financial support.DK is the recipient of an Australian Research Council Future Fellowship (project number FT160100344) funded by the Australian Government.

