## Supplementary Information for "A novel, ultrasensitive approach for quantitative carbohydrate composition and linkage analysis using LC-ESI ion trap tandem mass spectrometry"

#### ABSTRACT

Glycan identification and characterisation is essential to correlate glycoconjugate structure to biological function. The structural assignment of carbohydrates is often based on MS composition analyses and knowledge on well-studied glycosylation pathways. Nevertheless, many monosaccharide building blocks are indistinguishable by mass alone and detailed linkage information is also not easily obtained by MS/MS analyses, in particular when organisms are studied where the glycosylation pathways are less well defined. Here, we present a novel, simple and sensitive method using Reversed Phase (RP) – Liquid Chromatography Electrospray ionisation tandem mass spectrometry (LC–ESI-MS/MS) for unambiguous identification and linkage determination of monosaccharides including N-acetylneuraminic acids. Sequential permethylation and reductive amination steps are employed prior and after acid hydrolysis to enable separation and differentiation of the various monosaccharides and their respective linkage positions. The well-established, monosaccharide specific methylation patterns allowed for the identification of the various derivatised monosaccharide alditols based on their retention time and tandem mass spectrometry fingerprint. Absolute quantitation can also be accomplished by including a set of internal standards, thus simultaneously providing qualitative and quantitative information on the monosaccharide residues present.

### Supplementary information

***Supplementary Table S1:*** *MS operational parameters used in the present investigation*

| ***Ionization mode*** | ***Positive mode ESI for monosaccharides*** |
|---|---|
| *Capillary exit* | *1-1.3 kV* |
| *ICC* | *On* |
| *Maximum accumulation time* | *50 ms* |
| *Target* | *200,000 (MS/MS)* |
| *Scan range* | *50-500* m/z *(for MS and MS/MS)* |
| *Isolation window* | *2.0* m/z |
| *MS/MS fragmentation amplitude* | *60.0 %* |
| *Smart fragmentation option* | *On (start amplitude 30%—end amplitude 200%)* |

**Supplementary** ***Table 2:*** *List of N-glycans standards used to evaluate the developed linkage analysis method.*

| *Glycan Id* | *Glycan* | *Name* |
|---|---|---|
| 1 | 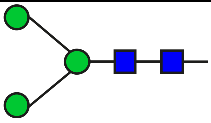 | M3 |
| 2 | 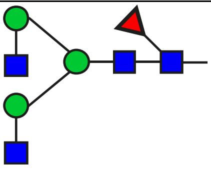 | GnGnF |
| 3 | 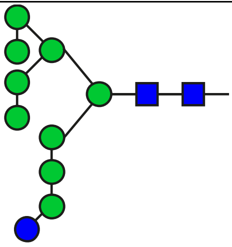 | M9 Glu1 |

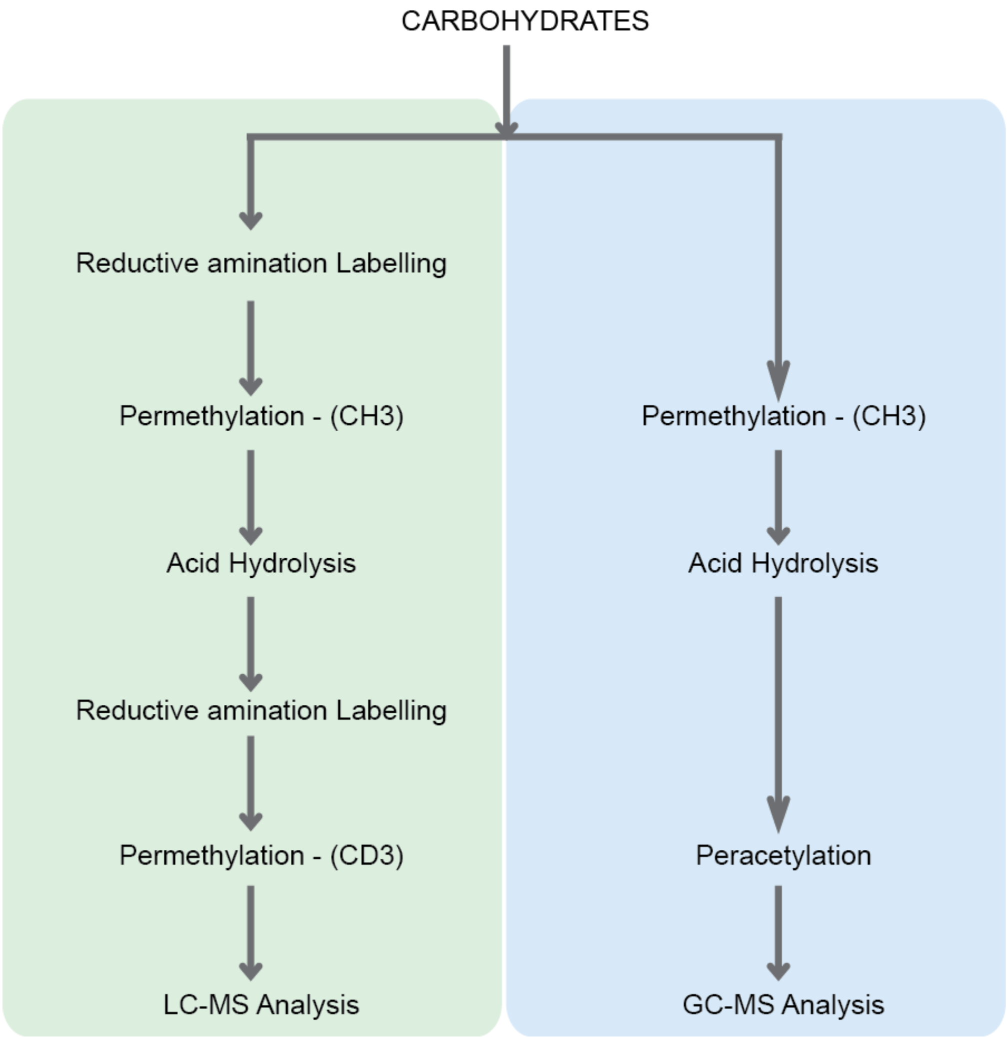


**Supplementary Figure 1***: Comparison of traditional GC-MS linkage analysis (right) versus the proposed strategy for linkage analysis using LC-MS developed in the course of this work (left). A detailed optimised step-by-step procedure for monosaccharide composition and linkage analysis using LC-MS/MS is provided in the supplementary methods section*.

### MATERIALS AND METHODS

#### OPTIMISED STEP-WISE PROCEDURE FOR MONOSACCHARIDE LINKAGE AND COMPOSITIONAL ANALYSIS

##### *Monosaccharide derivatization*

***Reducing end derivatization***

***Labelling Reagent 1:*** 0.35 M 2-Aminobenzamide and 1 M sodium cyanoborohydride were freshly prepared in dimethyl sulfoxide:acetic acid (7:3, v/v).

***Labelling Reagent 2:*** 0.35 M Aniline and 1 M sodium cyanoborohydride were freshly prepared in dimethyl sulfoxide:acetic acid (7:3, v/v).

1. Aliquot and freeze-dry the glycan sample in an Eppendorf tube
2. Add 50 µL of the labelling reagent 1 or 2.
3. Incubate for 120 min at 65°C.
4. Cool down to room temperature.
5. Add 500 µL of acetone or DCM and vortex for 30 s.
6. Centrifuge at 13,000 rpm for 5 min, the entire solvent was decanted and the procedure was repeated twice
7. Purify the derivatised carbohydrates using a carbon solid-phase extraction tip.

***$CH_3I$ permethylation***

8. Dry the glycan sample in glass tube and cap with a Teflon-lined screw cap.
9. Critical step: Samples must be dry and free of salts; otherwise, they will not completely methylate.
10. Add 500 µL of DMSO, cap and sonicate the tubes for 20–30 min.
11. Add approximately 50 mg of the powdered NaOH to each sample
12. Act quickly and do not attempt to weigh NaOH pellets, as they will absorb water rapidly.
13. Take care to use an even suspension of NaOH and to drop the reagent straight onto the sample—do not get the NaOH suspension on the side of the tube, or the reagent will be limiting.
14. Cap and sonicate for 20–50 min.
15. Add 30 µL of $CH_3I$ and sonicate for 10 min.

16. Caution: Only use $CH_3I$ in a fume hood, as it is extremely toxic and a suspected carcinogen.
17. Add 20 µL of $CH_3I$ and sonicate for 10 min.
18. Add 50 µL of $CH_3I$ and sonicate for 20 min.
19. Add 500 µL of water and 500 µL of DCM. Cap and vortex well (>40 s per sample). Centrifuge briefly to separate the phases.
20. Remove and discard the aqueous (upper) phase. Wash the lower DCM phase three times with 500 µL water.
21. Dry the lower DCM phase with a stream of dry nitrogen.

***C) Acid hydrolysis***

22. Add 200 µL of 2.0 M TFA to the methylated sample. Hydrolyse for 90 min at 121°C in a fan-forced oven.
23. Cool the sample and place the tube in a warm water bath (~30°C) and evaporate to dryness with a stream of nitrogen.
24. Alternatively samples can be transferred to an Eppendorf tube and dried in vacuo using a SpeedVac concentrator.
25. Add 100 µL of acetonitrile to the samples and dry the samples in the SpeedVac concentrator. Repeat this procedure three or five times to remove any residual TFA.
26. Purify the derivatised carbohydrates using a C18 solid-phase extraction tip.
27. Reductive amination derivatisation – 2AB

This is done exactly as described in the steps 1-4.

28. Derivatised samples can now be extracted via DCM: Water extraction. Lower DCM phase contains the derivatised glycan.

***D) $CD_3I$ Permethylation***

This is done exactly as described in the previous section 8-17 except that deuterated methyl iodide ($CD_3I$) is used instead.

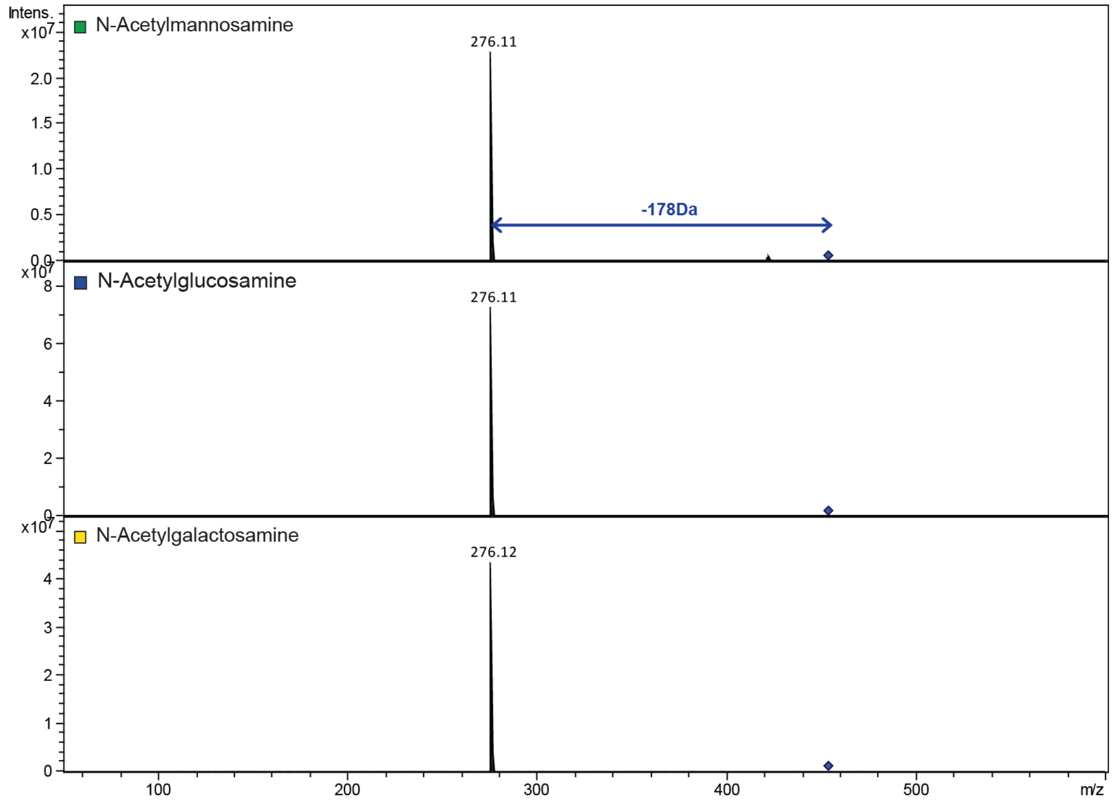


**Supplementary Figure 2***: Fragmentation pattern of 2-AB Me* N*-acetylhexosamines.*

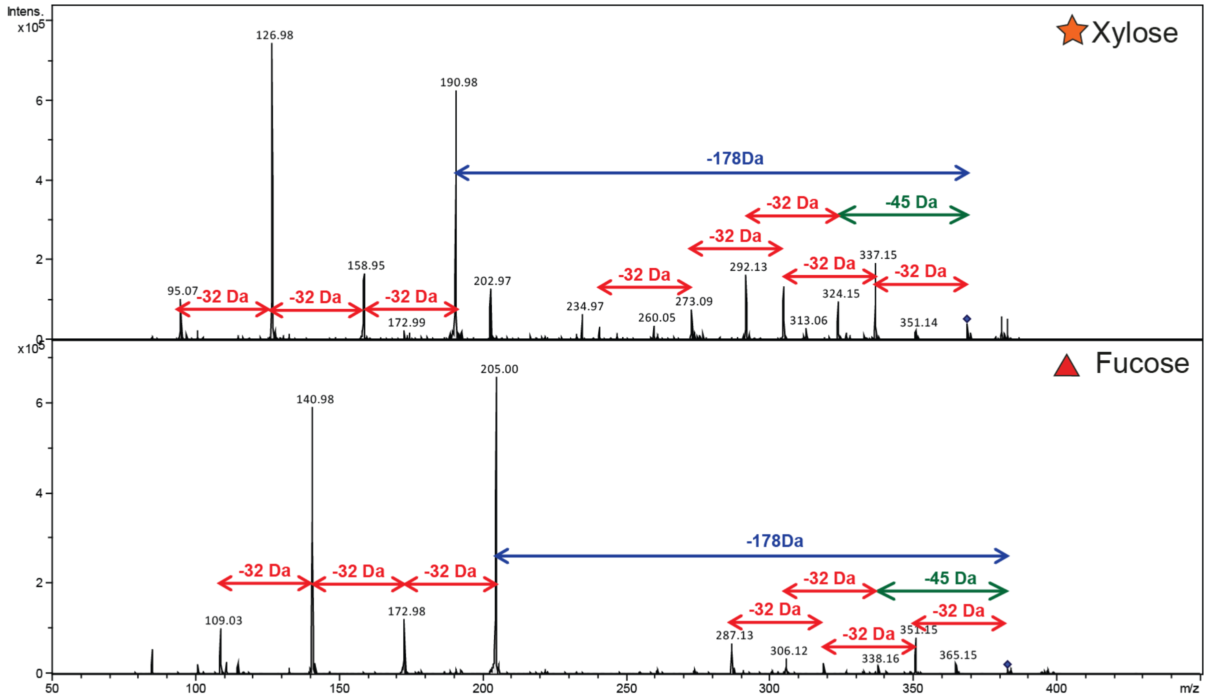


**Supplementary Figure 3***: Fragmentation pattern of 2-AB-Me xylose and fucose.*

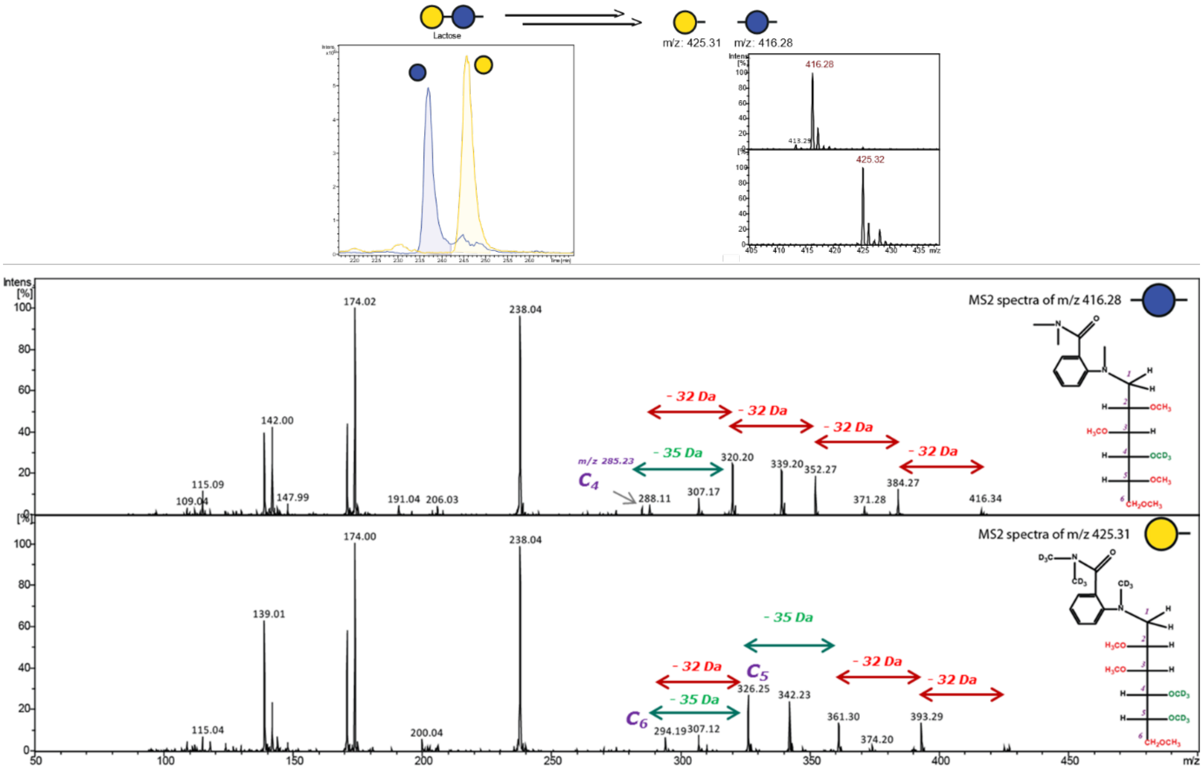


**Supplementary Figure** ***4:*** *LC-ESI-MS/MS analysis of 2-AB methylated glucose and galactose derived from lactose after linkage analysis.*

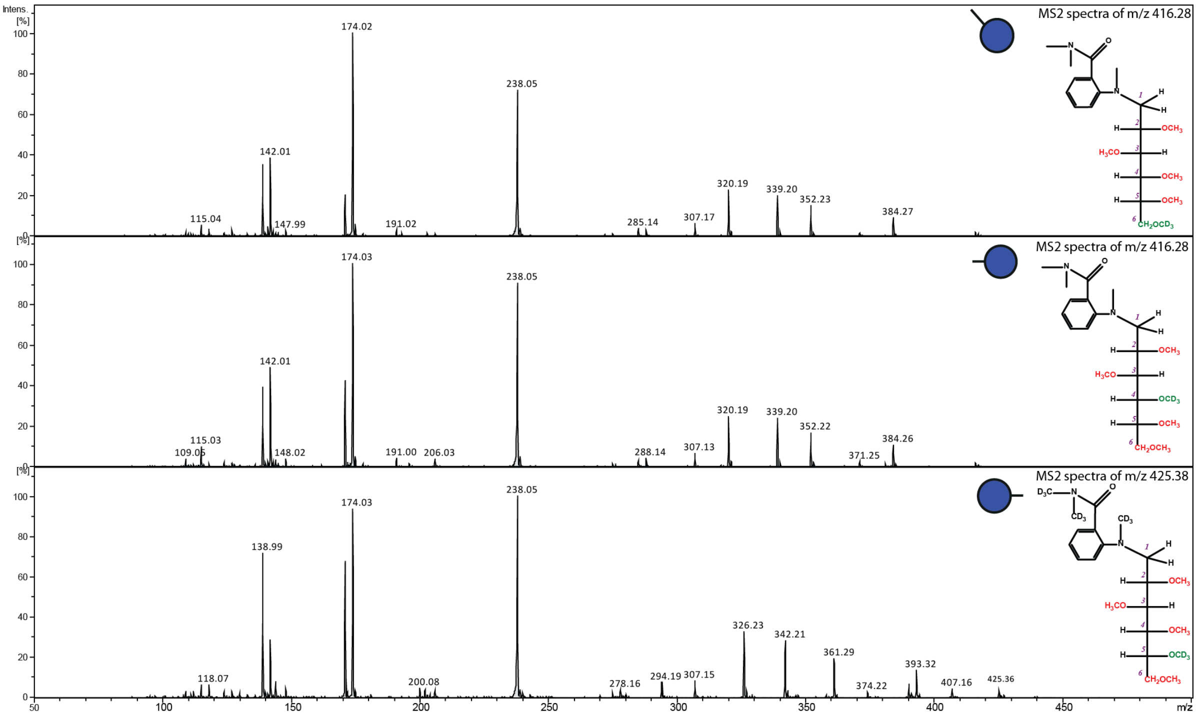


**Supplementary Figure** ***5: Comparison of 1-1 vs 1-4 vs 1-6 linked glucose residues derived from trehalose, lactose, and gentiobiose.*** *The linkage positions of the various glucose moieties were inferred from the sequential loss of either 32 Da or 35 Da from the precursor mass in the MS/MS spectra.*

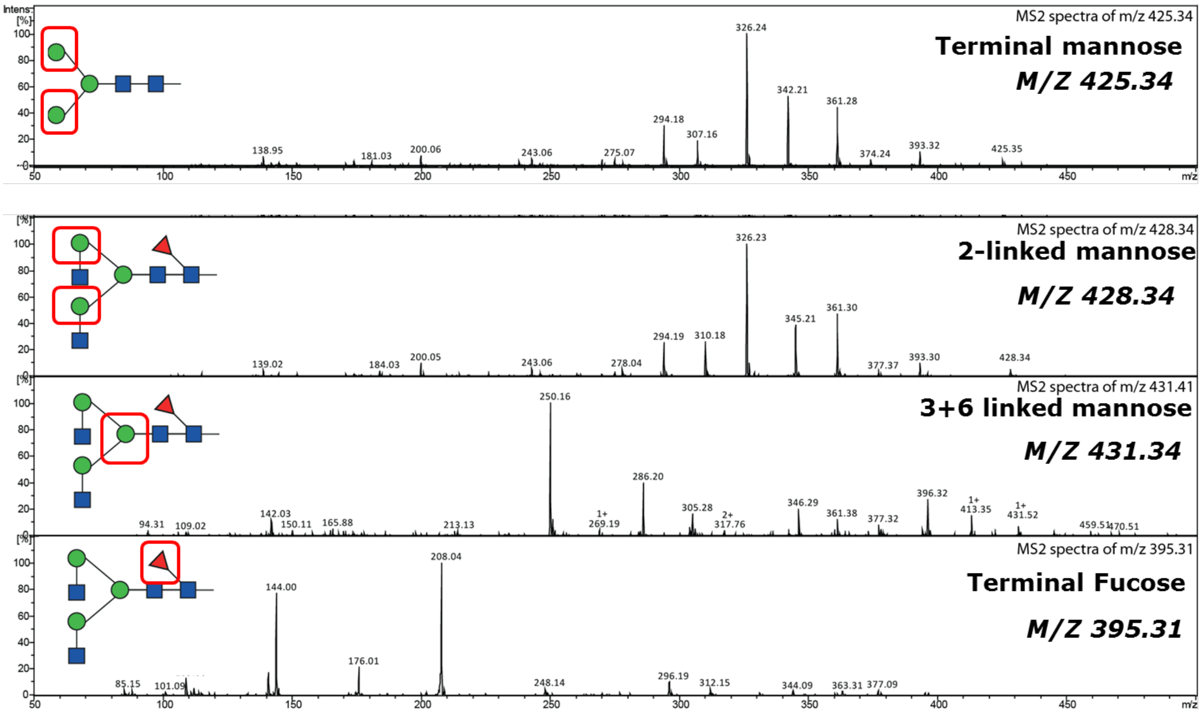


***Supplementary Figure 6:*** *Fragment spectra of various monosaccharide residues present in the standard N-glycans listed in table 2.*

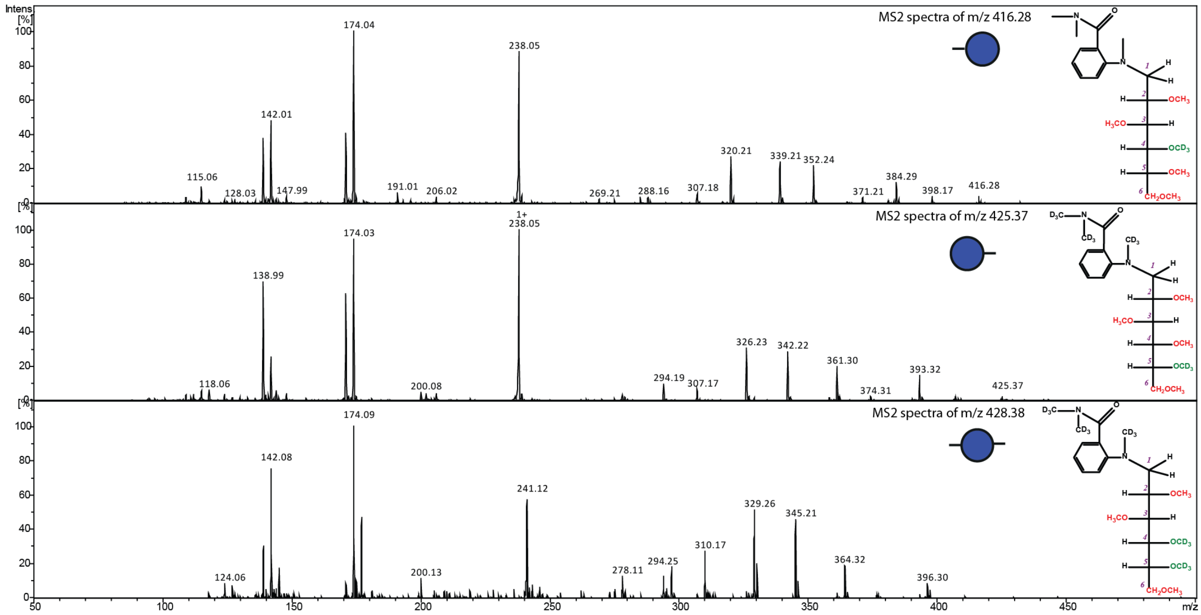


***Supplementary Figure 7:*** *Fragment spectra of reducing end, non-reducing end, and internal glucose residues present in the maltoheptaose. Linkage positions of the respective monosaccharides were identified based upon the molecular mass, the order of elution, and fragmentation spectra.*

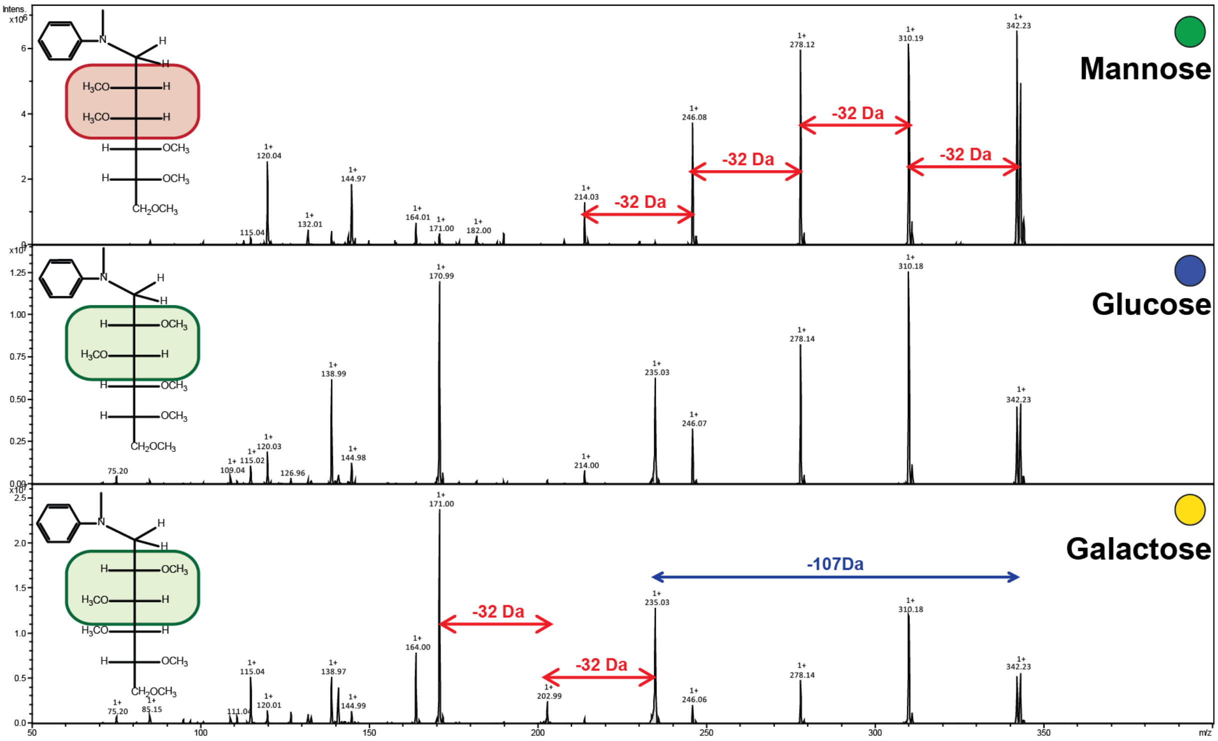


***Supplementary Figure 8:*** *Fragmentation pattern of various Aniline-Me hexoses of the protonated precursor ion* m/z *342.23. As described for 2-AB-Me hexose, the fragmentation behaviour depends upon the stereochemistry of the monosaccharide.*

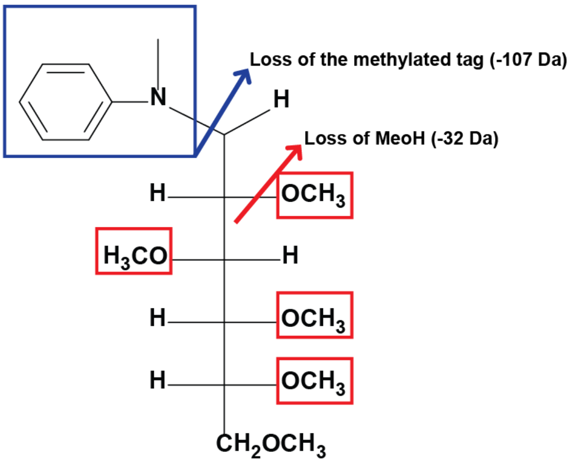


***Supplementary figure 9:*** *Fragmentation of Aniline-Me derivatized monosaccharides during electrospray ionization. Primary fragments occur between the C1 atom and the Nitrogen atom of the fluorescent tag, which results in the mass loss of 107 Da from the precursor. Secondary fragmentation results in the loss of methanol (32 Da) from the precursor mass.*

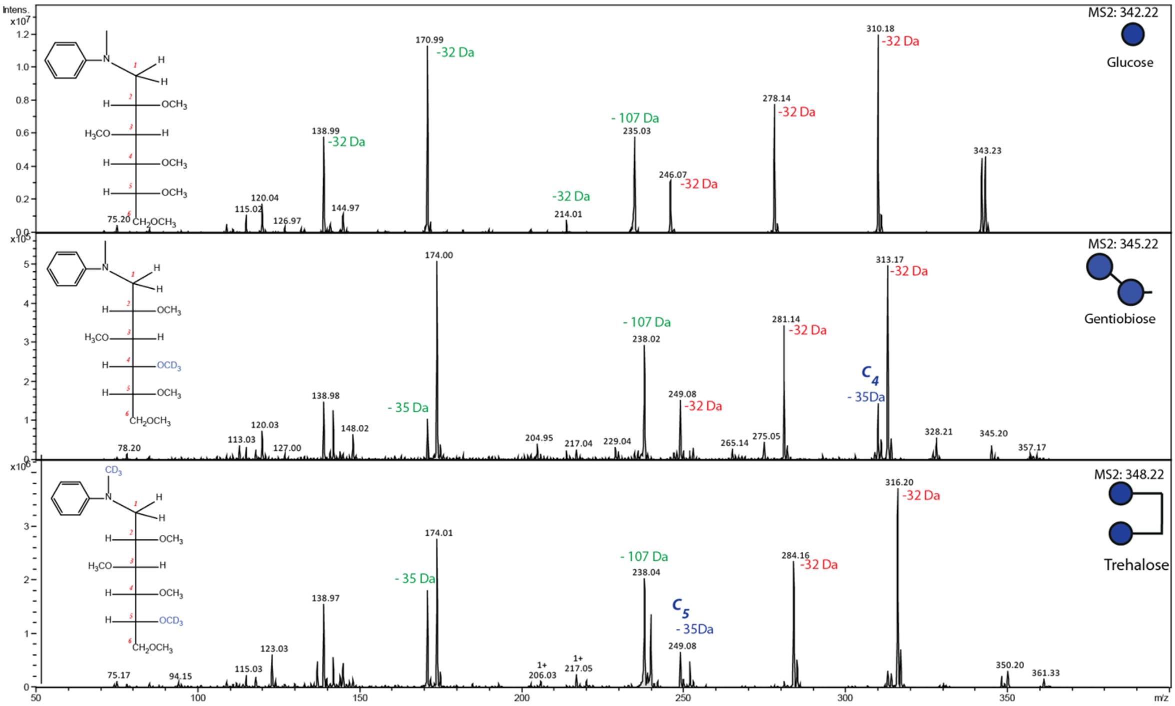


*Supplementary figure **10:** Linkage analysis using Aniline. Comparison of the fragmentation pattern of glucose vs 1-1 vs 1-4 linked glucose residues derived from gentiobiose and trehalose.*
